# SHIFT: speedy histological-to-immunofluorescent translation of whole slide images enabled by deep learning

**DOI:** 10.1101/730309

**Authors:** Erik A. Burlingame, Mary McDonnell, Geoffrey F. Schau, Guillaume Thibault, Christian Lanciault, Terry Morgan, Brett E. Johnson, Christopher Corless, Joe W. Gray, Young Hwan Chang

## Abstract

Spatially-resolved molecular profiling by immunostaining tissue sections is a key feature in cancer diagnosis, subtyping, and treatment, where it complements routine histopathological evaluation by clarifying tumor phenotypes. In this work, we present a deep learning-based method called speedy histological-to-immunofluorescent translation (SHIFT) which takes histologic images of hematoxylin and eosin-stained tissue as input, then in near-real time returns inferred virtual immunofluorescence (IF) images that accurately depict the underlying distribution of phenotypes without requiring immunostaining of the tissue being tested. We show that deep learning-extracted feature representations of histological images can guide representative sample selection, which improves SHIFT generalizability. SHIFT could serve as an efficient preliminary, auxiliary, or substitute for IF by delivering multiplexed virtual IF images for a fraction of the cost and in a fraction of the time required by nascent multiplexed imaging technologies.

**KEY POINTS:**

- Spatially-resolved molecular profiling is an essential complement to histopathological evaluation of cancer tissues.
- Information obtained by immunofluorescence imaging is encoded by features in histological images.
- SHIFT leverages previously unappreciated features in histological images to facilitate virtual immunofluorescence staining.
- Feature representations of images guide sample selection, improving model generalizability.

## INTRODUCTION

Physicians depend on histopathology—the visualization and pathological interpretation of tissue biopsies—to diagnose cancer. Hematoxylin and eosin (H&E)-stained histologic sections (3μm-thick formalin-fixed paraffin-embedded tissue biopsies) are the standard of care routinely employed by pathologists to make diagnoses. However, in challenging cases with indeterminate histology, or tumor differentiation, antibody labeling of tumor cells by a molecular imaging technique like immunofluorescence (IF) provides further characterization. It is becoming increasingly apparent that determining the spatially-resolved molecular profile of a cancer is important for disease subtyping and choosing a patient’s course of treatment (Duraiyan et al., 2012). Despite its clinical value, IF is time- and resource-intensive and requires expensive reagents and hardware, so assessment is typically limited to a small representative section of a tumor, which may not be fully representative of the neoplasm, which can be the case in areas of squamous differentiation in an adenocarcinoma (Hester et al., 2018). Also, the cost associated with IF may in some cases limit its use to within highly-developed clinical laboratories, further widening the quality-of-care gap between high- and low-income communities. The gaps between H&E and IF technologies highlights the broader need for automated tools that leverage information attained by a low-cost technology to infer information typically attained by a high-cost technology.

Recent advances in digital pathology and deep learning (DL) have made it possible to automatically extract valuable, human-imperceptible information from H&E-stained histology images (Campanella et al., 2019; Chen et al., 2018; Liu et al., 2017; Rivenson et al., 2019b, 2019a). Apart from histology images, Christiansen et al. and Ounkomol et al. each described supervised DL-based methods for inferring fluorescence images from transmitted light images of unlabeled human or rat cell lines or cell cultures, but not in complex, associated human tissues. Their methods were also based on strictly supervised learning frameworks, which are known to produce incoherent or discontinuous patterns in the virtual stains for some markers, though some of the authors suggest that an adversarial learning framework could address the problem (Christiansen et al., 2018). We previously introduced an adversarial DL-based method called speedy histological-to-IF translation (SHIFT) and demonstrated its ability to infer fluorescence images from images of H&E-stained tissue from a single patient with pancreatic ductal adenocarcinoma (PDAC) (Burlingame et al., 2018). In the current study, we test the generalizability of virtual IF staining by SHIFT through model validation on a limited but morphologically heterogeneous PDAC dataset comprised of samples from multiple patients.

DL models require a large amount of heterogeneous training data to generalize well across the population from which the training data was drawn. Since data limitations are common to many biomedical data domains (Campanella et al., 2019; Costa et al., 2018; Udrea and Mitra, 2017; Xiao et al., 2018), we begin by exploring the possibility that the choice of training samples can be optimized by selecting the few samples that are most representative of the population of samples at our disposal. Some DL-based applications have been proposed for histological image comparison and retrieval (Hegde et al., 2019; Otálora et al., 2018), but to the best of our knowledge none have been proposed for the express purpose of training set selection in a data-limited biomedical imaging domain. We describe the use of a data-driven DL-based method to select samples that optimizes the morphological heterogeneity of the dataset and promotes SHIFT model generalizability. As a proof of concept, we objectively measure the ability of SHIFT models to infer the spatial distribution of a pan-cytokeratin (panCK) antibody which marks cancer cells. We also show preliminary results for inference on the stromal marker *α*-smooth muscle actin (*α*-SMA), a first step toward the development of a generalized platform for multiplexed virtual IF imaging in human tissues. The advantages of a validated SHIFT model over traditional IF include (1) eliminating the need for expensive imaging hardware, bulk reagents, and technical undertaking of IF protocols; (2) virtual IF images can be generated in near-real time; and (3) the portability of SHIFT allows it to be integrated into existing imaging workflows with minimal effort.

## RESULTS

### Building a Dataset of Spatially-registered H&E and IF Images

SHIFT requires spatially-registered pairs of H&E and IF whole slide images (WSIs) for model training and testing (**Figure 1A**). Such data would usually be acquired by processing two adjacent tissue sections, one stained by H&E and another stained by IF, then spatially registering the images into the same coordinate system based on their shared features (Chang et al., 2017). Unfortunately, this can lead to inconsistencies between H&E and IF image contents when high-frequency cellular features differ between adjacent sections, even when the sections are as few as 5μm apart. To alleviate this issue, we developed a protocol that allows for H&E and IF staining in the same section of tissue (see **METHODS**). Clinical samples of PDAC from four patients (Samples A, B, C and D) were chosen via pathological review of archival H&E images as exemplifying a spectrum of both histological differentiation and heterogeneity (**Figure 1B**). Chosen samples were sectioned, processed, and stained with DAPI nuclear stain and panCK monoclonal antibody; the staining was confirmed and the slides were scanned. After scanning, the coverslips were removed and the slides were stained with the designed modified H&E protocol, permanently cover slipped and then scanned again. Nuclear information from the hematoxylin and DAPI stains in pairs of H&E and IF images were used to register images in a common coordinate system. Images were then pre-processed to minimize noise and account for technical variability in staining and image acquisition. To exclude regions of autofluorescence that greatly diminished the signal-to-noise ratio of the real IF images, images from samples B and D were subdivided into image subsets {B1, B2, B3} and {D1, D2, D3, D4, D5}.

**Figure 1.**
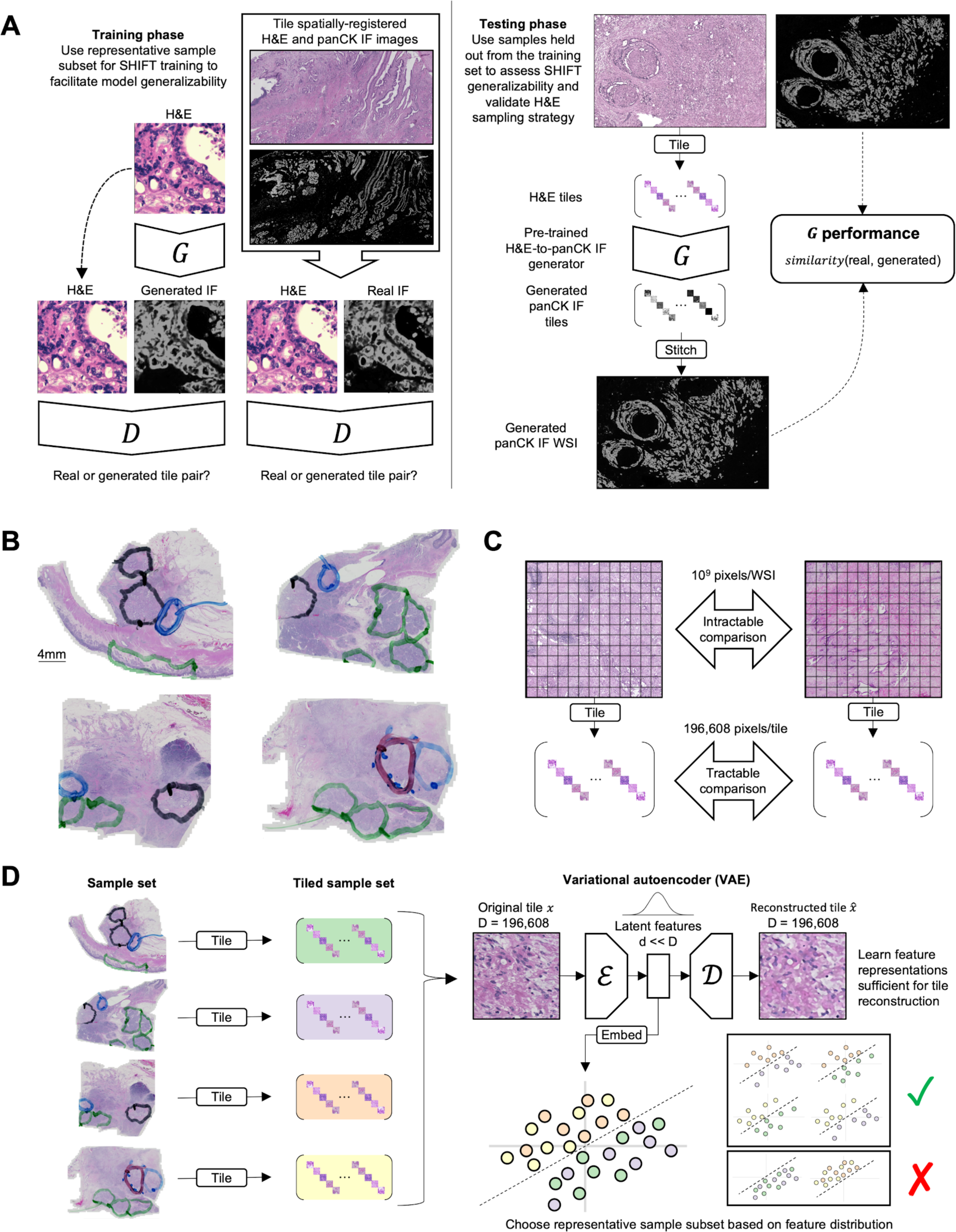
Overview of virtual IF staining with SHIFT and feature-guided H&E sample selection. (A) Schematic of SHIFT modeling for training and testing phases. The generator network *G* generates virtual IF tiles conditioned on H&E tiles. The discriminator network *D* learns to discriminate between real and generated image pairs. See also **Figure S2**. (B) Four heterogeneous samples of H&E-stained PDAC biopsy tissue used in the current study. Pathologist annotations indicate regions that are benign (green), grade 1 PDAC (black), grade 2/3 PDAC (blue), and grade 2/3 adenosquamous (red). (C) Making direct comparisons between H&E whole slide images (WSIs) is intractable because each WSI can contain billions of pixels. By decomposing WSIs into sets of non-overlapping 256×256 pixel tiles, we can make tractable comparisons between the feature-wise distribution of tile sets. (D) Schematic of feature-guided H&E sample selection. First, H&E samples are decomposed into 256×256 pixel tiles. Second, all H&E tiles are used to train a variational autoencoder (VAE) to learn feature representations for all tiles; for each 196,608-pixel H&E tile in the dataset, the encoder *ε* learns a compact but expressive feature representation that maximizes the ability of the decoder 𝒟 to reconstruct the original tile from its feature representation (see **METHODS**). Third, the tile feature representations are used to determine which samples are most representative of the whole dataset.

### Feature-guided Identification of Representative Histological Samples

For a SHIFT model to generalize well across the population of PDAC samples, it must be trained on a representative subset of the population, which motivated the development of a means to quantitatively compare images (see **METHODS**). In particular, we wished to learn which sample—or sequence of samples—should be selected to build a training set that is most representative of the population of samples. As a consequence of their large dimensions, direct comparison between gigapixel H&E images is intractable, so we decomposed each image into sets of non-overlapping 256×256 pixel tiles (**Figure 1C**). Even the small 256×256 pixel H&E tiles contain 196,608 (256×256×3 channels = 196,608) pixel values each and are difficult to compare directly. To establish a more compact but still expressive representation of the H&E tiles, we trained a variational autoencoder (VAE) (Kingma and Welling, 2013)--an unsupervised DL-based method for representation learning and feature extraction--to learn 16-dimensional feature representations of each tile (see **METHODS**), which makes comparing tiles more tractable (**Figure 1D**). Using the dimensionality reduction algorithm *t*-SNE (van der Maaten and Hinton, 2008), we embed all the H&E tiles into a 2D map to visualize the learned feature space over which the H&E tiles are distributed, which provides a visual aid to our selection of the most representative set of samples (**Figures 1D**).

Using the VAE which we pre-trained on all samples, we extracted features from each H&E tile (**Table S1)** and assessed how each feature was distributed across samples, finding that several of the features discriminated between samples (**Figure 2A**). In particular, the bimodal distribution of some features suggested two sample clusters, one formed by samples A and B, and another formed by samples C and D. These clusters were corroborated by visualization of the sample tiles embedded in the reduced-dimension feature space generated by *t*-SNE (**Figure 2B** and **Table S1**). Since the feature distributions of the H&E tiles highlighted the redundancy between clustered samples, we reasoned that a balanced selection of samples from each cluster would yield a more representative training set and ultimately improve SHIFT model generalizability. Using the full 16-dimensional feature representations of the H&E tiles and an information-theoretic framework for representative sample selection (Feng Pan et al., 2005; see **METHODS**), we were able to quantitatively identify sample B and the duo of samples B and D as the single and two most representative samples, respectively, and were then considered for training sets in subsequent experiments. **Figure S1** illustrates the feature distributions of several sample combinations in comparison to that of the full dataset.

**Figure 2.**
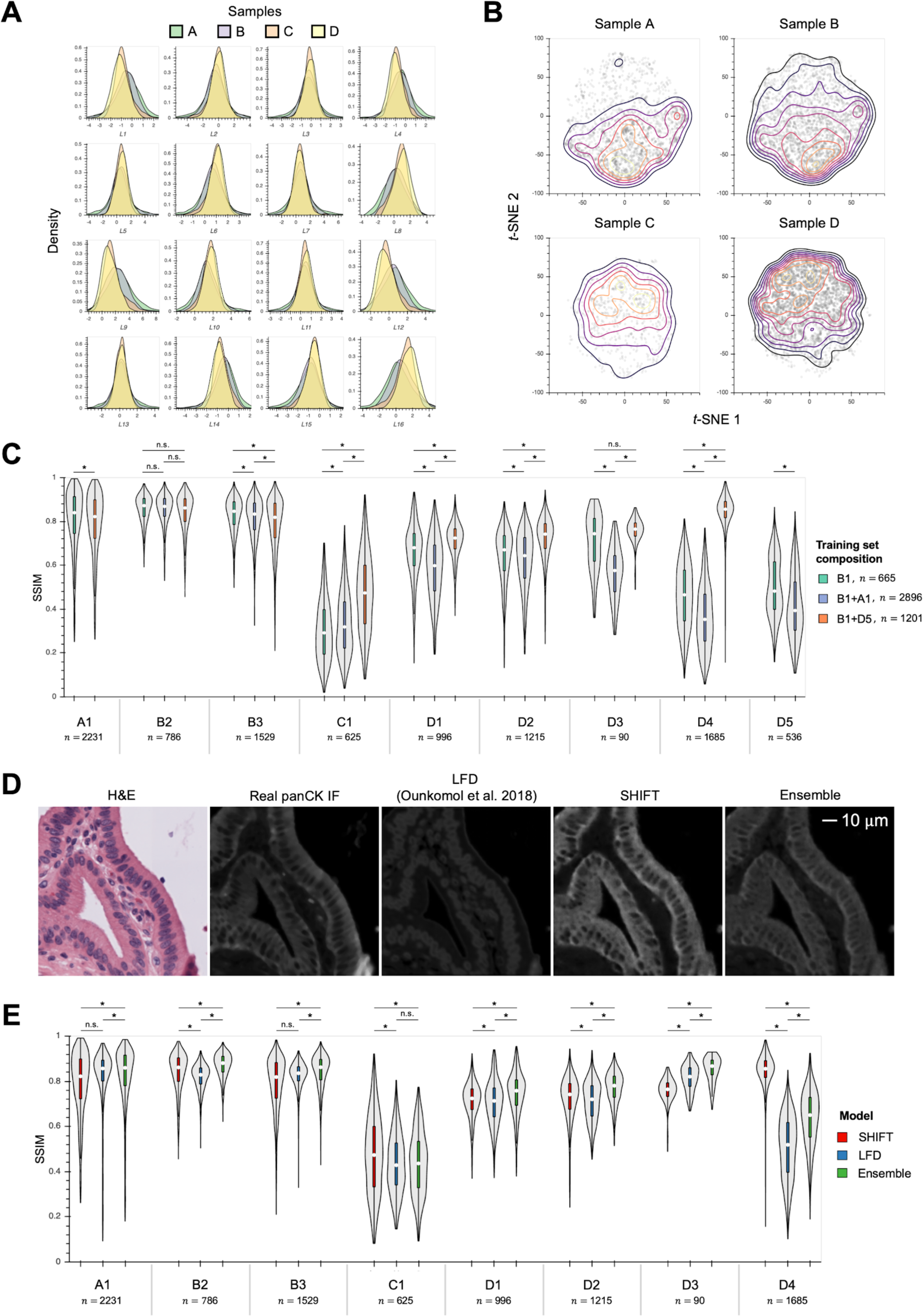
Feature-guided H&E sample selection and virtual IF staining with SHIFT. (A) Distribution of the 16 latent features (L1-L16) extracted by VAE from sample H&E tiles. (B) *t*-SNE embedding of latent feature representations of sample H&E tiles, faceted by sample identity. Each point in each plot represents a single H&E tile. Contour lines indicate point density. (C) SHIFT model test performance for optimal and non-optimal training set sample compositions. The paired H&E and IF images from samples B and D were subdivided into smaller images B = {B1,B2} and D = {D1,D2,D3,D4,D5} to avoid regions of IF that exhibited substantial autofluorescence. The x-axis labels indicate sample identity, where each letter corresponds to a unique sample and each number corresponds to a subset of that sample. Each *n* denotes the number of image tiles that were extracted from that sample. Plots for sample subsets are not show if that sample subset was a component of a model’s training set. *p<.05; for three group comparisons we used the Friedman test with Nemenyi post-hoc test; for two group comparisons we used the Wilcoxon signed-rank test. White dots in violin plots represent distributional medians. (D) Visual comparison of virtual staining methods. The ensemble results are attained by averaging the output images of SHIFT and Label-Free Determination models. See also **Figure S2**. (E) Test performance comparison of virtual staining methods. The x-axis labels indicate sample identity, where each letter corresponds to a unique sample and each number corresponds to a subset of that sample. Each *n* denotes the number of image tiles that were extracted from that sample. Plots for sample subsets B1 and D5 are not show because those sample subsets were components of the models’ training sets. *p<.05; Friedman test with Nemenyi post-hoc test. White dots in violin plots represent distributional medians.

### Virtual IF Staining in Histological Samples

SHIFT models are built on an adversarial image-to-image translation framework (Isola et al., 2016), with a regularization strategy designed to improve inference on sparse IF images (**Figures 1A and S2**; see **METHODS**). Adversarial learning frameworks compute their losses over images, in contrast to strictly supervised learning frameworks where losses are computed over pixels, which is likely to improve model inference in many virtual staining tasks (Christiansen et al., 2018). Having identified the most representative samples in our dataset, we next tested whether or not a SHIFT model could learn a correspondence between H&E and IF images that generalizes across samples.

We hypothesized that if tissue and cell morphologies observed in H&E-stained tissue are a function of a given marker, then it should be possible to infer the spatial distribution of that marker based on the H&E-stained tissue alone; that is, H&E-to-IF translation should be learnable and generalizable such that a model can be extended to samples from patients that were not included in the training set. To test this hypothesis, we trained SHIFT models to generate virtual IF images of the cancer marker panCK conditioned on input H&E images alone (**Figure 1A**). To simultaneously assess the utility of our sample selection method, we trained models using different combinations of sample subsets in the training set. Training sets consisted of paired H&E and IF image tiles from either sample subset B1 (from the most representative single sample), sample subsets B1 and D5 (from the most representative duo of samples), or sample subsets A1 and B1 (from a less representative duo of samples as counterexample). Sample subsets B1 and D5 were selected because they contained a similar number of tiles, providing a balance between the sample clusters. Model performance was quantified by measuring the structural similarity (SSIM) (Rivenson et al., 2019b; Wang et al., 2004), a widely used measure of image similarity as perceived by the human visual system, between corresponding virtual and real IF images from samples left out of the model’s training set. The SSIM between two images is calculated over pixel neighborhoods in the images and provides a more coherent measure of image similarity than pixel-wise measures like Pearson’s *r*. Considering the performance of the model trained on B1 as the baseline, we see a marked improvement in model generalizability across samples when the more representative sample subsets B1 and D5 are used for training than when the less representative sample subsets A1 and B1 are used (**Figure 2C**). By stitching together the virtual IF tiles from a given sample in the test set, we were able to make large-scale comparisons between real and virtual panCK IF images (**Figure S3**). We also experimented with SHIFT inference of the stromal marker *α*-SMA (**Figure S4**).

SHIFT is not the only virtual staining method to have been recently proposed. Label-Free Determination (LFD) (Ounkomol et al., 2018) is a supervised DL-based virtual staining method which produces models that were shown to have learned the relationship between images of cell cultures visualized by transmitted light or fluorescence, where sub-cellular structures have been labeled with genetically-encoded fluorescent tags. Because the SHIFT generator *G* and the LFD are both based on the popular U-net architecture (Ronneberger et al., 2015), we compared these models that generate images using a similar architecture, but have differing training formulae and loss functions (see **METHODS**). To make a fair comparison between the adversarial SHIFT and supervised LFD models, we trained a LFD using the representative sample subsets B1 and D5, matching the number of optimization steps taken by the SHIFT model that was trained using the same training set (**Figure S5**). In addition to the performance of independent SHIFT and LFD models, we also considered the ensemble result, taken as the average image of the SHIFT and LFD output images. Across all samples in the test set, either SHIFT alone or the ensemble of SHIFT and LFD tended to perform better than LFD alone (**Figure 2C**).

## DISCUSSION

Spatially-resolved molecular profiling of cancer tissues by technologies like IF provides more information than routine H&E histology alone. However, the rich information obtained from IF comes at significant expense in time and resources, restricting IF access and use. Here, we present and extend the validation of SHIFT, a DL-based method which takes standard H&E-stained histology images as input and returns virtual IF images of inferred biomarker distributions. Using a limited but heterogeneous dataset, we demonstrated that SHIFT models are able to generalize across samples drawn from different PDAC patients, even for training sets that are over an order of magnitude smaller than the test set (train*n* = 665and test *n* = 9693for models trained on sample subset B1 only). Results from our sampling experiments are consistent with the expectation that an automated and quantitative method for representative sample selection will be critical to the effective development and deployment of DL models on large-scale digital pathology datasets. Finally, we compared the adversarial SHIFT method with an alternative, supervised virtual staining method and found that the virtual staining task tends to be best accomplished by the ensemble of both methods. Based on the success of DL-based ensemble methods in other biomedical domains (Codella et al., 2017; Xiao et al., 2018), we expect ensemble methods to become increasingly relevant to the development of virtual staining applications.

While we have demonstrated the utility of SHIFT for only a limited number of virtual stains, there are emerging opportunities to test the extensibility of our approach. Following recent advances in multiplexed immunofluorescence and immunohistochemistry (mIF/IHC) technology (Goltsev et al., 2018; Lin et al., 2016; Reiß et al., 2019; Tsujikawa et al., 2017), it is now possible to visualize tens or hundreds of distinct biomarkers in a single tissue section. On their own, these technologies promise a more personalized medicine through a more granular definition of disease subtypes, and will undoubtedly broaden our understanding of cellular heterogeneity and interaction within the tumor microenvironment, both of which play increasingly important roles in the development and selection of effective treatments (Lu et al., 2019; Yuan, 2016). With a paired H&E and mIF/IHC dataset that encompasses the expression of hundreds of markers within serial tissue sections, we could begin to quantify the mutual information between histology and expression of any biomarker of interest.

Though our results suggest broad applicability, there are limitations and challenges to both the feature-guided sampling and virtual staining methods we present here. Both methods assume a relationship between H&E and IF representations of tissue, so they may fail to maximize representativeness and make incorrect inference in the IF domain if there is no association between histology and a given biomarker. Even when there is an association, finding a meaningful way to compare real and virtual images remains a challenge, as we have experienced in experiments modeling biomarkers with fine, high-frequency distributions like *α*-SMA (**Figure S4**). Like other virtual staining methods that have been deployed on whole human tissues (Rivenson et al., 2019b, 2019a), we used SSIM as a measure of image similarity between real and virtual IF images. This classical perceptual measure is used in many imaging domains, but we found that it is sensitive to perturbations commonly associated with image registration and technical or instrumentation noise (**Figure S6**). In light of this, we advocate for the development and use of perceptual measures that are more aware of such perturbations and better correlate with human perception of image similarity or quality (Azulay and Weiss, 2018; Patel et al., 2019). It must be restated that our results are supported by a dataset comprised of samples from just four patients, so some fluctuation in performance between samples should be expected, and indeed was observed (**Figure S3C**). Our goal in choosing a relatively small dataset was to demonstrate that, even when limited, SHIFT could learn a general relationship between H&E and IF tissue representations, and we believe that the fluctuation in performance between samples could be addressed by increasing the sample size. With the emergence of digital pathology datasets containing tens of thousands of whole slide images (Campanella et al., 2019), the opportunities to improve virtual staining technologies are only becoming more numerous.

Since SHIFT can infer virtual IF images as H&E-stained tissue section are imaged, SHIFT could provide pathologists with near-real-time interpretations based on standard H&E-stained tissue. Therefore, SHIFT could serve as an efficient preliminary, auxiliary, or substitute technology for traditional IF in both research and clinical settings by delivering comparable virtual IF images for a fraction of the cost and in a fraction of the time required by traditional IF or mIF/IHC imaging. As such, we see SHIFT as an opportunity to simultaneously economize and democratize advanced imaging technologies in histopathology workflows, with implications for multiplexed virtual imaging. Further, we see our method of optimal selection of representative histological images, which promotes morphological heterogeneity in the training set, as a complement to data augmentation, transfer learning, and other means of addressing the problem of limited training data. Moreover, this method will contribute to saving resources and minimizing unnecessary efforts to acquire additional staining or manual annotation for DL applications in biomedical imaging.

## AUTHOR CONTRIBUTIONS

Conceptualization, E.A.B. and Y.H.C.; Methodology, E.A.B, M.M., B.E.J., and Y.H.C.; Software, E.A.B., G.F.S., G.T., and Y.H.C.; Validation, E.A.B., C.L., T.M., and C.C.; Formal Analysis, E.A.B., C.L., T.M., and Y.H.C.; Investigation, E.A.B., M.M., C.L., T.M., B.E.J., C.C., and Y.H.C.; Resources, E.A.B., B.E.J., J.W.G., and Y.H.C.; Data Curation, E.A.B., M.M., and Y.H.C.; Writing--Original Draft, E.A.B., M.M., G.F.S., T.M., and Y.H.C.; Writing--Review & Editing, E.A.B., M.M., G.F.S., G.T., T.M., B.E.J., C.C., J.W.G. and Y.H.C.; Visualization, E.A.B and M.M.; Supervision, B.E.J., C.C., J.W.G. and Y.H.C.; Project Administration, E.A.B., M.M., B.E.J., and Y.H.C.; Funding Acquisition, E.A.B., G.F.S., Y.H.C., B.E.J., and J.W.G.

## ACKNOWLEDGMENTS

This work was supported in part by the National Cancer Institute (U54CA209988, U2CCA233280), the OHSU Center for Spatial Systems Biomedicine, the Brenden-Colson Center for Pancreatic Care, and a Biomedical Innovation Program Award from the Oregon Clinical & Translational Research Institute. We acknowledge expert technical assistance by staff in the Advanced Multiscale Microscopy Shared Resource, supported by the OHSU Knight Cancer Institute (NIH P30 CA069533) and the Office of the Senior Vice President for Research. Equipment purchases included support by the OHSU Center for Spatial Systems Biomedicine, the MJ Murdock Charitable Trust, and the Collins Foundation. We acknowledge the Histopathology Shared Resource for pathology studies, which is supported in part by the University Shared Resource Program at Oregon Health and Sciences University and the Knight Cancer Institute (P30 CA069533, and P30 CA069533 13S5). The resources of the Exacloud high performance computing environment developed jointly by OHSU and Intel and the technical support of the OHSU Advanced Computing Center are gratefully acknowledged. E.A.B. receives support from a scholar award provided by the ARCS Foundation Oregon. B.E.J. was supported by a Postdoctoral Fellowship, PF-17-031-01-DDC, from the American Cancer Society.

## METHODS

### Resources

| REAGENT or RESOURCE | SOURCE | IDENTIFIER |
| --- | --- | --- |
| **Antibodies** |  |  |
| Alpha-smooth muscle actin monoclonal antibody (1A4 (ASM-1)) | ThermoFisher Scientific | https://www.thermofisher.com/antibody/product/Alpha-Smooth-Muscle-Actin-Antibody-clone-1A4-asm-1-Monoclonal/MA5-11547 |
| Goat anti-mouse IgG2a cross-adsorbed secondary antibody, AlexaFluor® 555 | ThermoFisher Scientific | https://www.thermofisher.com/antibody/product/Goat-anti-Mouse-IgG2a-Cross-Adsorbed-Secondary-Antibody-Polyclonal/A-21137 |
| Pan-cytokeratin monoclonal antibody (AE1/AE3), AlexaFluor® 488, EBioscience | ThermoFisher Scientific | https://www.thermofisher.com/antibody/product/Pan-Cytokeratin-Antibody-clone-AE1-AE3-Monoclonal/53-9003-82 |
| Ki-67 (D3B5) monoclonal antibody, AlexaFluor® 647, Conjugate #12075 | Cell Signaling Technology | https://www.cellsignal.com/products/antibody-conjugates/ki-67-d3b5-rabbit-mab-alexa-fluor-647-conjugate/12075?N=4294960176&Nrp=100&fromPage=plp |
| **Biological Samples** |  |  |
| Human PDAC tumor samples | Oregon Pancreas Tissue Registry | IRB: 3609 |
| **Chemicals, Peptides, and Recombinant Proteins** |  |  |
| SlowFade Gold Antifade Mountant with DAPI | ThermoFisher Scientific | https://www.thermofisher.com/order/catalog/product/S36938 |
| Normal goat serum blocking solution | Vector laboratories | https://vectorlabs.com/normal-goat-serum-blocking-solution.html |
| **Deposited Data** |  |  |
| Imaging data | This manuscript | https://gitlab.com/eburling/shift/ |
| **Software and Algorithms** |  |  |
| Python | Python Software Foundation | RRID:SCR_008394 |
| PyTorch | Paszke et al., 2017 | https://pytorch.org/ |
| imgaug | Jung, 2018 | https://github.com/aleju/imgaug |
| Netron | https://www.lutzroeder.com/ai/ | https://github.com/lutzroeder/netron |
| HoloViews | Stevens et al., 2015 | https://holoviews.org/ |
| Bokeh | Bokeh Development Team, 2019 | https://bokeh.pydata.org/ |
| scikit-image | Walt et al., 2014 | https://scikit-image.org/ |
| scikit-learn | Pedregosa et al., 2011 | https://scikit-learn.org |
| SciPy | Jones et al., 2001 | https://www.scipy.org/ |
| Jupyter | Kluyver et al., 2016 | https://jupyter.org/ |

### Human tissue samples

This work was conducted as part of the Oregon Pancreas Tissue Registry (IRB#3609), which was approved by the Institutional Review Board at Oregon Health and Science University (OHSU). Informed consent was obtained from all subjects. Four cases of pancreatic ductal adenocarcinoma (PDAC) diagnosed as moderately differentiated adenocarcinoma were retrieved from the OHSU Surgical Pathology Department. Sample A was from a male aged 83 at diagnosis; sample B was from a female aged 74 at diagnosis; sample C was from a female aged 57 at diagnosis; and sample D was from a female aged 73 at diagnosis. H&E-stained sections were secondarily reviewed by two board-certified surgical pathologists tasked to identify and classify areas of tumor heterogeneity in representative sections from each case. Discrepancies between pathologists were ameliorated by consensus review. Samples were chosen via pathological review as exemplifying a spectrum of both histological differentiation and morphology.

### Pathological evaluation of human tissue samples

Gold standard review of histologic sections by pathologists tasked with identifying heterogeneous differences in PDAC tumor morphology and grade revealed interobserver agreement in the identification of areas of squamous differentiation in one case and various tumor grades within neoplasms in the other three cases. All four cases were predominantly grade 2 adenocarcinoma and there was no disagreement evaluating marked regions of interest. The case with areas of squamous differentiation did not clearly meet the 30% threshold for adenosquamous classification. The other three cases were predominantly grade 2 with foci of grade 1 and others with grade 3.

### Immunofluorescence staining

#### Preparation of tissue for immunofluorescence staining

Formalin-fixed paraffin-embedded tissue blocks were serially sectioned by the OHSU Histopathology Shared Resource. From each block, three sections were cut in order to generate a standard H&E for pathological review and downstream analysis, a second serial section of tissue for immunofluorescence staining/post-immunofluorescence H&E staining, and a third section for secondary only control. After sectioning, the second serial tissue section was immediately baked at 55°C for 12 hours and subjected to standard deparaffinization; the slides underwent standard antigen retrieval processing, washing, and blocking. Upon completion, primary antibodies were diluted and applied.

#### Application of antibodies

Alpha-Smooth Muscle Actin (Mouse monoclonal antibody, IgG2a, Clone: 1A4; Pierce/Invitrogen, cat#MA5-11547) was diluted to 1:200 with Ki-67 (D3B5), (Rabbit monoclonal antibody, IgG, Alexa Fluor® 647 Conjugate; Cell Signaling Technology, cat#12075S) diluted to 1:400, along with Pan Cytokeratin (AE1/AE3) (Mouse monoclonal antibody, IgG1, Alexa Fluor® 488 Conjugate; ThermoFisher, cat#53-9003-82), which was diluted to 1:200 in 10% Normal Goat Serum in 1% Bovine Serum Albumin in Phosphate Buffered Saline. Primary antibodies were diluted and incubated overnight at 4°C. After incubation, secondary antibody (Goat anti-mouse monoclonal antibody, IgG2A, Alexa Fluor® 555 Conjugate; Life Technologies, cat# A21137), at 1:200 dilution was applied to the slides and incubated at room temperature for one hour. After incubation slides were washed and mounted with Slowfade Gold Antifade Mountant with DAPI (Fisher Scientific, cat#S36936) in preparation for image acquisition.

#### Post-IF H&E staining of tissue samples

After the IF stained slides were scanned and the immunofluorescence staining verified, the glass coverslips were removed and the slides were immediately processed for post-IF H&E staining. Post-IF H&E staining was performed with the Leica Autostainer XL staining system at the OHSU Histopathology Shared Resource with the modified staining protocol described in the table below:

|  |  |
| --- | --- |
| Hematoxylin | 10 minutes |
| Wash in water | 1 minute |
| Acid alcohol (.5% HCl in 70% Ethanol) | 8 seconds |
| Wash in water | 25 seconds |
| Bluing solution | 2 minutes |
| Wash in water | 20 seconds |
| 80% Ethanol/water | 25 seconds |
| Eosin | 10 seconds |
| 80% Ethanol/water | 25 seconds |
| 95% Ethanol/water | 20 seconds |
| 100% Ethanol (two times) | 25 seconds |
| Xylene (five times) | 25 seconds |

#### Image acquisition and presentation

Slides were scanned with the Zeiss Axio Scan.Z1 slide scanner of the OHSU Advanced Multiscale Microscopy Shared Resource with the 20X objective in both brightfield and immunofluorescence scanning. Carl Zeiss Images (CZI) were acquired using Zeiss Zen software. CZI images from the Zeiss Axioscan Slide Scanner were processed with the Zeiss Blue Zen Lite microscope software package. All brightfield and immunofluorescence images were manually annotated and exported as TIFF files for downstream image processing.

### Image pre-processing

Raw H&E and IF whole slide images (WSIs) must be pre-processed to remove technical noise, account for between-sample intensity variation, and align paired H&E and IF WSIs in a shared coordinate system. To do so, we use the following pipeline:

1. Downscaling: 20X WSI are downscaled by a factor of 2 in x and y dimensions to generate 10X WSIs. We experimented with using either 20X or 10X images and found that models performed best when using 10X images.
2. Registration: H&E and IF WSIs are spatially registered using SURF features (Bay et al., 2006; Chang et al., 2017) extracted from hematoxylin and DAPI binary masks of nuclei generated by Otsu’s method, respectively.
3. Technical noise reduction: IF WSIs are median filtered with a 5-pixel-radius disk structuring element.
4. Intensity normalization: H&E WSI pixel intensities are normalized as previously described (Macenko et al., 2009). Following Christiansen et al., IF WSI pixel intensities are normalized to have a fixed mean=0.25 and standard deviation=0.125, then clipped to fall within [0,1].
5. Image tiling: WSIs are tiled into non-overlapping 256×256 pixel tiles, such that each H&E tile has a corresponding spatially-registered IF tile. H&E tiles that contained more than 50% background were removed along with the corresponding IF tiles. Each 10X WSI is comprised of hundreds or thousands of non-overlapping 256×256 pixel tiles.

### Feature-guided training set selection

Although DL approaches like SHIFT and Label-Free Determination require substantial training data to be robust and generalizable, due to resource constraints we hope that a small number of paired H&E and IF image samples is required for model training. Typically, archival WSIs of H&E-stained tissue sections exist on-hand for each sample, which allows for the screening of samples to identify the minimal number of samples that maximally represent the morphological spectrum of the disease being considered. The recent works of Hegde et al. and Otálora et al. demonstrate that DL systems are well-suited for image retrieval tasks in digital pathology, wherein a pathologist submits a query image or region of interest and the DL system returns similar images based on their DL-defined feature representations. We seek to solve the inverse task of heterogeneous training set selection in digital pathology, though our approach could be extended to any data-limited biomedical imaging domain.

Since PDAC is a morphologically heterogeneous disease, building a representative training set is crucial to the design of a model that will generalize across heterogeneous biopsy samples after deployment. In order to minimize the required resources for acquiring paired H&E and IF images but still cover a broad spectrum of heterogeneous morphological features in the selected H&E samples, we propose a clustering method to learn a heterogeneous representation of H&E sample images. To assess the morphological features of each sample, we use a variational autoencoder (VAE) (Kingma and Welling, 2013) to extract 16-dimensional feature vectors from each H&E tile to establish comparisons between samples. Since texture and morphological features on H&E tiles in each cluster of samples will be comparatively more similar than those of the other cluster, we only select representative H&E samples from each cluster for our training dataset. We also tried using other feature vector sizes for representation learning, e.g. 2, 4, 8, 32, but found that a feature vector size of 16 yielded the lowest reconstruction losses.

For example, if there are four samples being considered for IF staining, but resources limit the number of samples that can be stained to two, a decision must be made about which samples should be selected. For the four samples, we aggregate their archival H&E WSIs, extract features from H&E tiles for each sample using a VAE, and quantitatively determine the samples needed to maximally cover the feature space over which the H&E tile set is distributed. By screening and selecting samples in this data-driven fashion, we exclude homogeneous or redundant samples that would not contribute to model generalizability. This maximizes model performance by ensuring that our training dataset is representative of the disease being modeled, thus minimizing cost through the selection of the fewest samples required to do so. Perhaps more importantly, when we fail to generate reliable virtual IF images for certain tissue samples or IF markers, this framework will be useful to examine whether or not their morphological features are well presented in the training dataset, which can guide how we select additional samples when updating our dataset.

To identify the sequence of samples that should be selected, we adapt a sample selection algorithm (Feng Pan et al., 2005) parameterized using the following notation:

| Parameter | Description |
| --- | --- |
| $X$ | Complete tile set of all samples, $X = \{x_1, x_2, \dots, x_n\}$ |
| $x_i$ | Single tile, $x_i \in X$ |
| $X_i$ | Subset of $X$ corresponding to the $i$th sample, $X_i \subset X$ |
| $F$ | Complete VAE-learned feature set, $F = \{f_1, f_2, \dots, f_m\}$ |
| $f_i$ | Single feature, $f_i \in F$ |
| $A$ | Random variable defined over $F$ |
| $T$ | Random variable defined over $X$ |

We begin with a tiles × features table, where we set *m* = 16 for our experiments:

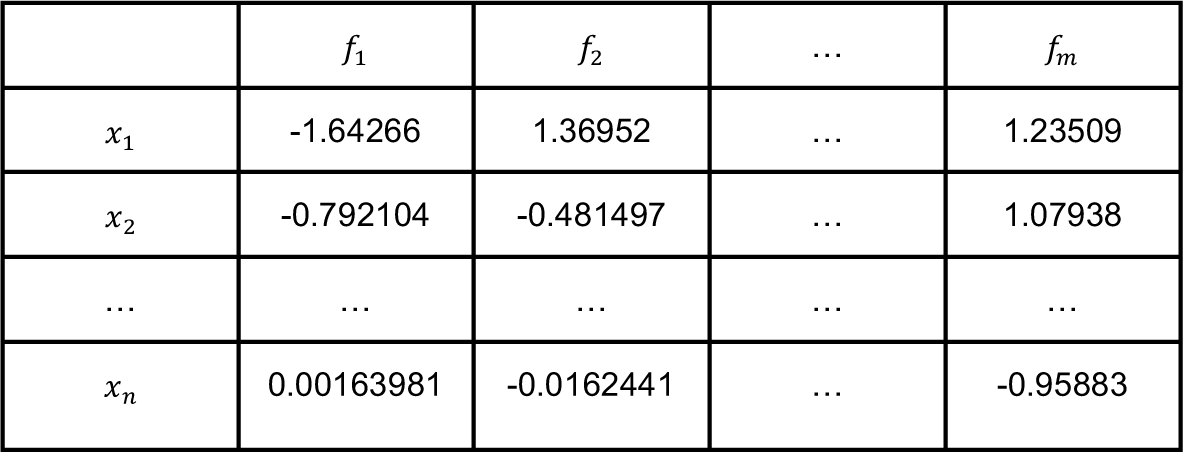

We normalize across rows of the table, such that each tile is now represented as a probability distribution over the feature domain:

| | $f_1$ | $f_2$ | ... | $f_m$ | Sum |
| --- | --- | --- | --- | --- | --- |
| $x_1$ | 0.00418311 | 0.0850498 | ... | 0.0743519 | 1 |
| $x_2$ | 0.0148384 | 0.0202433 | ... | 0.0964193 | 1 |
| ... | ... | ... | ... | ... | 1 |
| $x_n$ | 0.0208721 | 0.0205021 | ... | 0.00798802 | 1 |

We define the random variables *T*and *A*over tile domain *X*and the feature domain *F*, respectively, such that *P*(*A* = *f*_1_|*T* = *x*_1_) = 0.418311, *P*(*A* = *f*_2_|*T* = *x*_2_) = 0.0202433, and so on. With this conditional probability table, we can define probability distributions for each subset 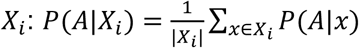 To measure the representativeness of sample *X*_6_to the full dataset *X*, we compute the Kullback-Leibler (KL) divergence between*P*(*A*|*X*_*i*_) and 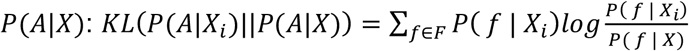 We then weight this divergence by the proportion of *X* that *X*_*i*_ comprises, 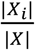, to prioritize subsets that contribute many tiles to *X*. We define the single most representative sample as 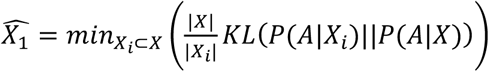, the most representative duo of samples as 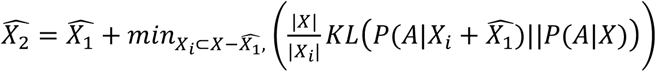, the most representative trio of samples as 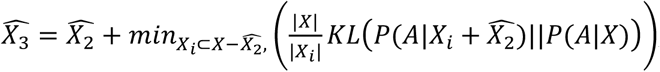, and so on. In this way, we define the sequence of samples that should be chosen to optimally increase the representativeness of the training set.

### Model architectures

#### Conditional generative adversarial networks

Image-to-image translation—the mapping of pixels from one scene representation to pixels of another representation of the same scene—is a fundamental image processing problem. Conditional generative adversarial networks (cGANs) (Goodfellow et al., 2014; Mirza and Osindero, 2014) are a compelling deep learning-based solution to the image-to-image translation problem which have been deployed for many tasks, including detection of skin lesions (Udrea and Mitra, 2017), retinal image synthesis (Costa et al., 2018), super-resolution fluorescence image reconstruction (Ouyang et al., 2018), and virtual H&E staining (Rivenson et al., 2019b). To approach the problem of translating H&E images to their IF counterparts, SHIFT adopts the cGAN-driven architecture *pix2pix* (Isola et al., 2016), which benefits from its bipartite formulation of generator and discriminator. Like other methods proposed for image-to-image translation, cGANs learn a functional mapping from input images *x* to ground truth target images *ŷ* but, unique to a cGAN architecture, it is the task of a generator network *G* to generate images *ŷ* conditioned on *x*, i.e. *G*(*x*) = *ŷ* that fool an adversarial discriminator network *D*, which is in turn trained to tell the difference between real and generated images. What ensues from this two-network duel is a *G* that generates realistic images that are difficult to distinguish from real images, some GAN-generated images being sufficiently realistic to be considered as a proxy for the ground truth when labeled data are scarce or prohibitively expensive (Bousmalis et al., 2016). Concretely, the cGAN objective is posed as a binary cross-entropy loss:

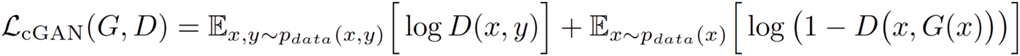

where *G* seeks to minimize the objective and thus minimize the distinguishability of generated and real images, while *D* seeks the opposite. In addition to the task of fooling *D, G* is also encouraged to generate images that resemble real images through incorporation of an L1 reconstruction loss term:

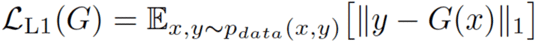

The full cGAN objective is:

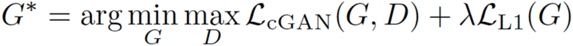

where the L1 tuning parameter *λ* is adapted according to the IF stain prevalence in the current batch of IF tiles (Burlingame et al., 2018). Training data consist of spatially registered pairs of H&E image tiles (*x*) and IF image tiles (*y*), while the test data consist of H&E and IF image pairs withheld from the training data. Models were trained using the Adam optimizer with a learning rate of 0.002 for 500 epochs. Training batch sizes were set to 64. The first layers of both the generator and discriminator networks were 128 filters deep (see **Figure S2** for additional architectural details). Full model details are available at https://gitlab.com/eburling/shift/.

#### Variational autoencoders

Variational autoencoder (VAE) architecture (Kingma and Wellington, 2013) is designed to elucidate salient features of data in a data-driven and unsupervised manner. A VAE model seeks to train a pair of complementary networks: an encoder network θ that seeks to model an input *x*_*i*_ as a hidden latent representation *z*_*i*_, and a decoder network ϕ that seeks to reconstitute *x*_*i*_ from its latent representation *z*_*i*_. The VAE cost function shown below penalizes model training with an additional Kullback-Leibler (KL) divergence term that works to conform the distribution of *z* with respect to a given prior, which in our case is the standard normal distribution:

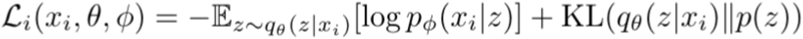

Where

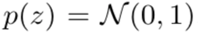

By specifying a latent dimension *z* less than the input dimension of *x*, a VAE model learns a pair of optimal encoding and decoding functions that enable reconstruction of an input sample subject to capacity constraints of the latent feature space within the model. In general, this formulation learns encoding functions that compress the information content in the high-dimensional input into a low-dimensional embedding space that learns dataset features sufficient to reconstitute the original input sample while preserving an expected distribution over the learned features. This interpretation enables a specified selection criteria function designed to sample whole slide images whose constituent tiles maximally cover the entire learned feature space with a minimal number of samples.

#### Model ensembles

In addition to testing the ability of independent SHIFT and LFD models to generate virtual IF images, we also tested model ensembles. Ensemble images were generated by simply averaging the virtual IF image outputs of SHIFT and LFD models trained to generate the same stain using the same training set.

### Imaging data augmentation

To boost the effective number of images in our training sets and improve model robustness against expected types of technical noise, we apply image augmentations to each image in each training batch using the Python library imgaug (Jung, 2018). We apply Gaussian blur, flipping, affine geometric transformation, Gaussian noise, Poisson noise, rotation, and add to hue and saturation in each channel. The implementation of our imaging data augmentation can be viewed at https://gitlab.com/eburling/shift.

### Image processing in figures

The IF images in Figure 1 are contrast enhanced by saturating the top 1% and bottom 1% of pixel intensities. All other images are processed as described in the image pre-processing section above.

### Image comparisons

To compare real and virtual IF images, we measure their structural similarity (Zhou Wang et al., 2004) using the compare_ssim function implemented in the Python library scikit-learn (Pedregosa et al., 2011). We calculate the SSIM between 11-pixel windows of the real and virtual IF image tiles. The SSIM between two windows *x* and *y* is defined as:

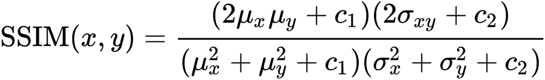

where *μ*_*x*_ and *μ*_*y*_ are the mean intensities of *x* and *y, σ*_*xy*_ is the covariance of *x* and *y*, 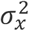 and 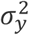 are the variances of *x* and *y*, and *c*_1_and *c*_2_ are stabilizing constants. The SSIM between real and virtual IF images is the SSIM averaged over their windows. Based on our observation that SSIM is sensitive to simulations of technical noise which are impossible for SHIFT models to infer (**Figure S6**), we apply Gaussian filtering to real and virtual IF images tiles before calculating SSIM using the gaussian function implemented in the Python library scikit-image with sigma set to 3. We also measured the Pearson’s correlation coefficient between images in **Figure S6** using the pearson_r function implemented in the Python library SciPy (Jones et al., 2001).

### Code availability

Code for training and inference can be found at https://gitlab.com/eburling/shift.

## SUPPLEMENTARY MATERIALS

**Table S1. Variational autoencoder-learned representations and t-SNE coordinates for H&E tiles.**

**Figure S1.**
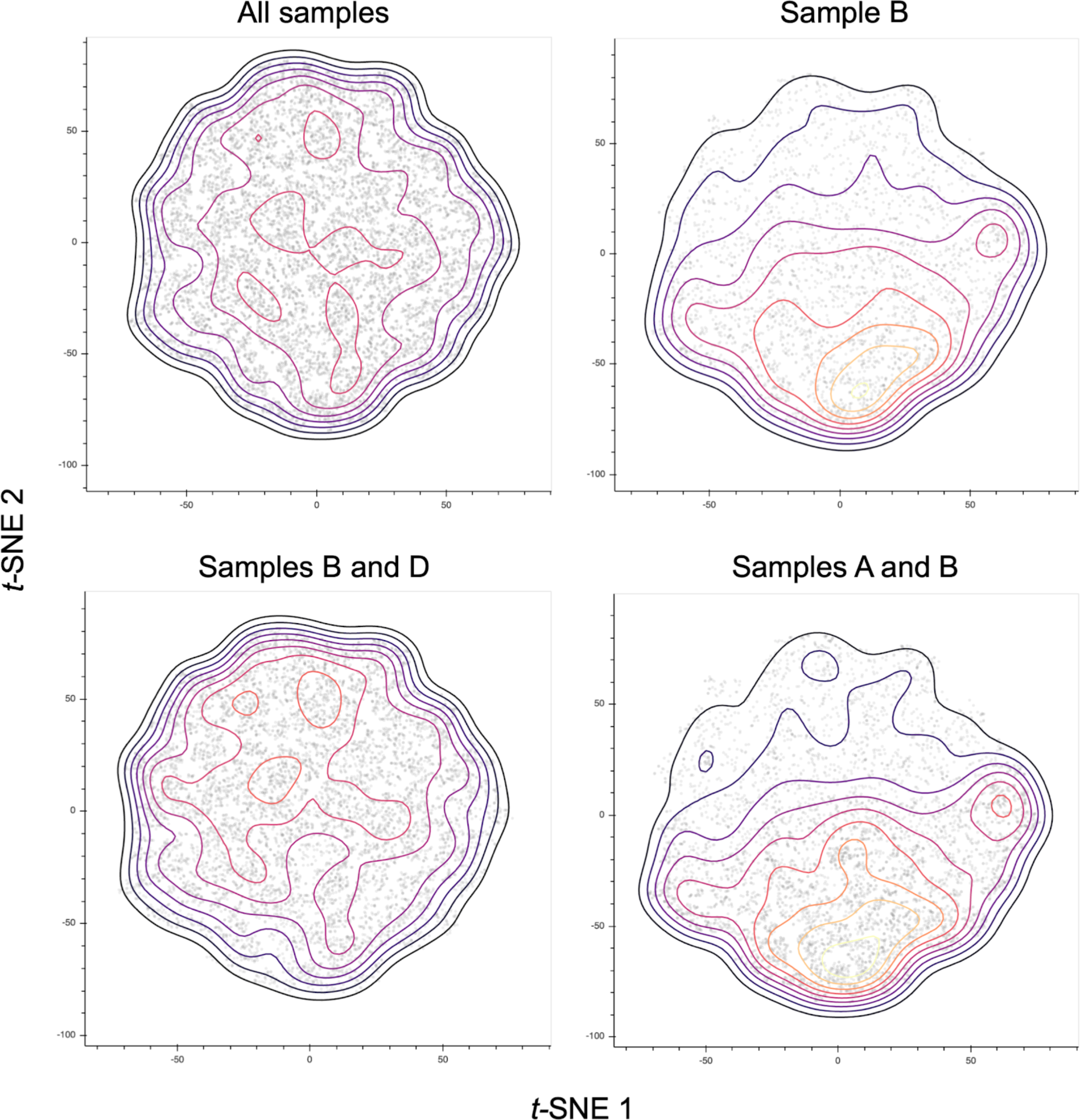
H&E tile feature distributions of experiment sample combinations. The VAE-learned 16-dimensional feature vector representations for each H&E are embedded into 2 dimensions using *t*-SNE. Each point in each plot represents a single H&E tile. Contour lines indicate point density. Sample B is the single sample that is most representative of the whole dataset. Samples B and D are the duo of samples that are most representative of the whole dataset. Samples A and B are a duo of samples that poorly represent the whole dataset.

**Figure S2.**
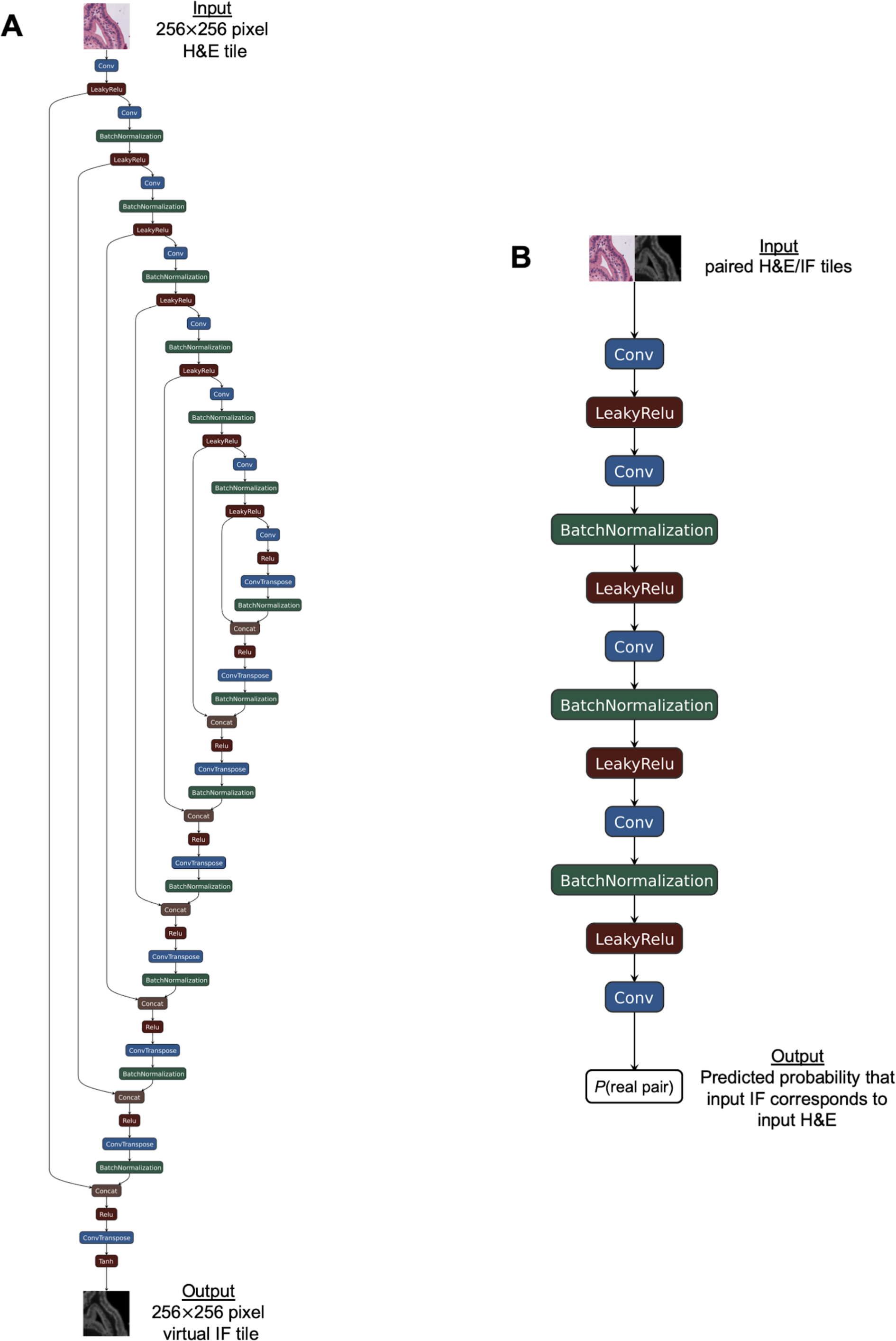
Schematics of cGAN architecture used by SHIFT. The cGAN architecture used by SHIFT is based on the *pix2pix* framework (Isola et al., 2016). Schematics were generated with the Netron network viewer tool (https://github.com/lutzroeder/netron). Full implementation details available on GitLab (https://gitlab.com/eburling/shift). (A) Architecture of generator network *G* which is based on the U-net architecture (Ronneberger et al., 2015). (B) Architecture of discriminator network *D*.

**Figure S3.**
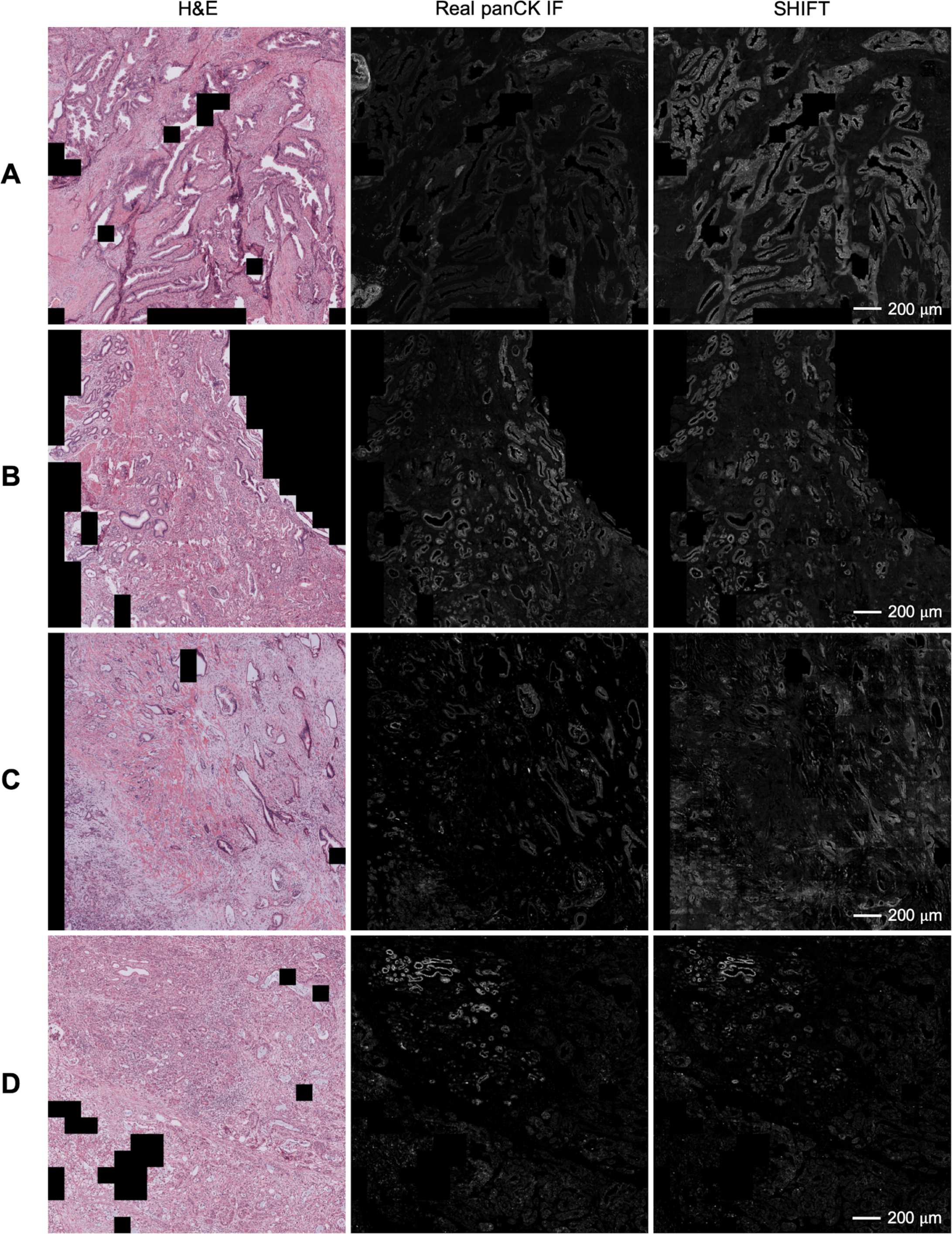
Large-scale comparison of real and virtual panCK staining generated by SHIFT. SHIFT images were generated by a model trained on sample subsets B1 and D5. Results show are from the test set. Tiles were excluded if they contained more than 50% background in the H&E representation (black tiles). (A) Representative images taken from sample A. (B) Representative images taken from sample B. (C) Representative images taken from sample C. (D) Representative images taken from sample D.

**Figure S4.**
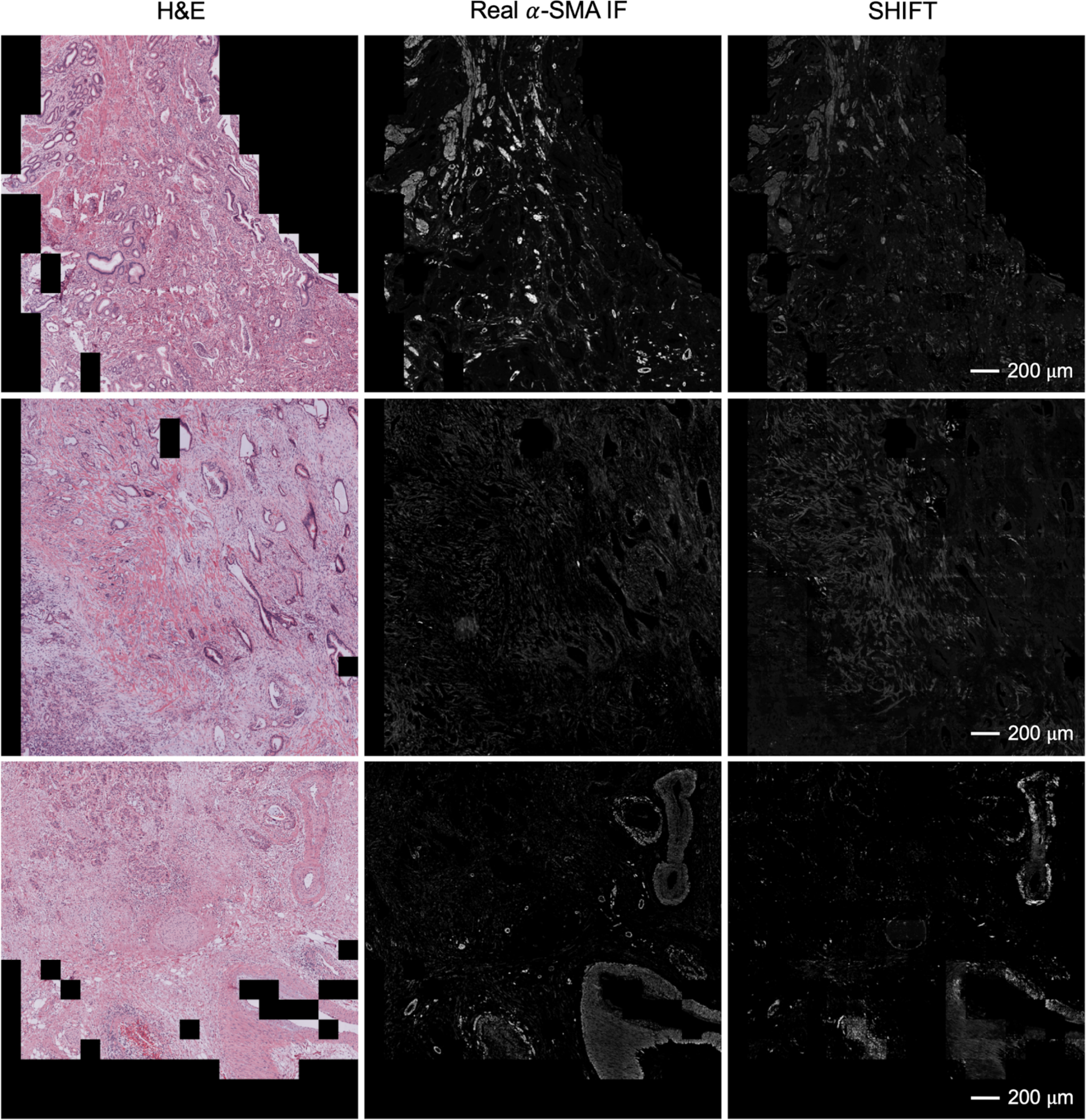
Large-scale comparison of real and virtual *α*-SMA staining generated by SHIFT. SHIFT images were generated by a model trained on sample subsets B1 and D5. Results show are from the test set. Tiles were excluded if they contained more than 50% background in the H&E representation (black tiles).

**Figure S5.**
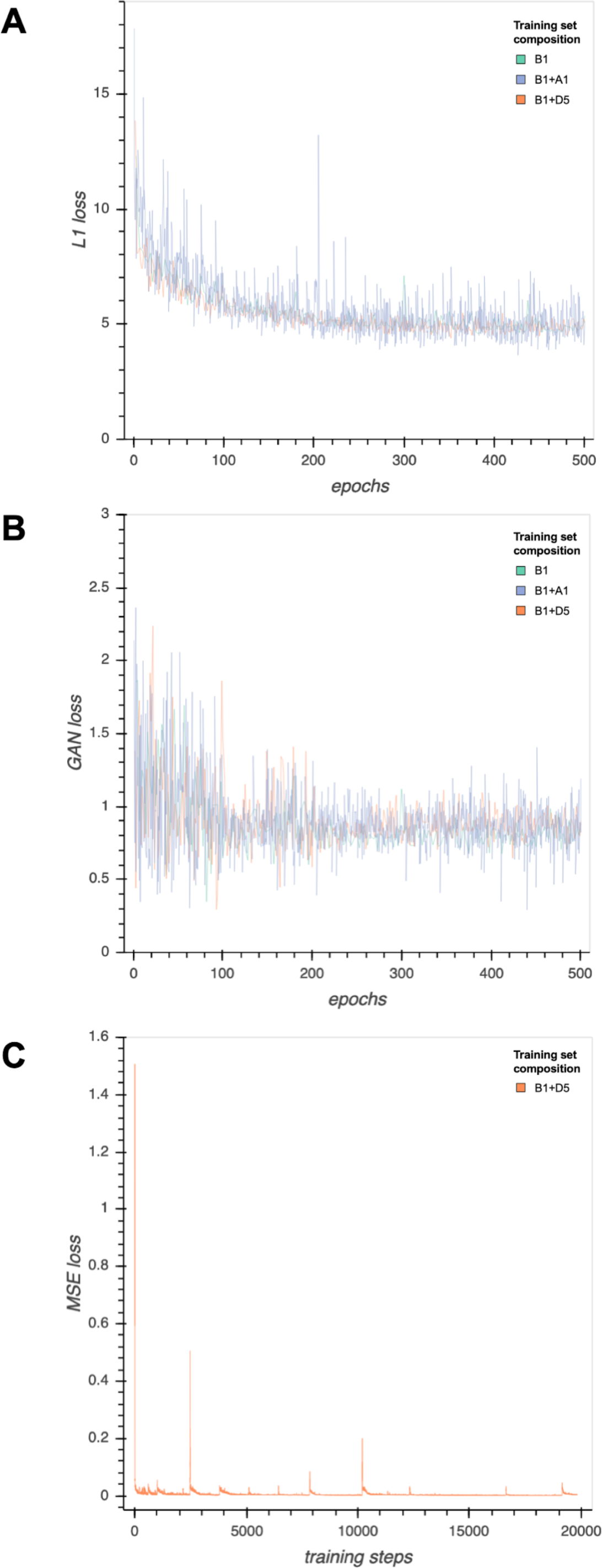
Training losses for virtual staining models. (A) L1 training loss for SHIFT models for each training set composition. (B) GAN training loss for SHIFT models for each training set composition. (C) Mean squared error (MSE) training loss for Label-Free Determination model.

**Figure S6.**
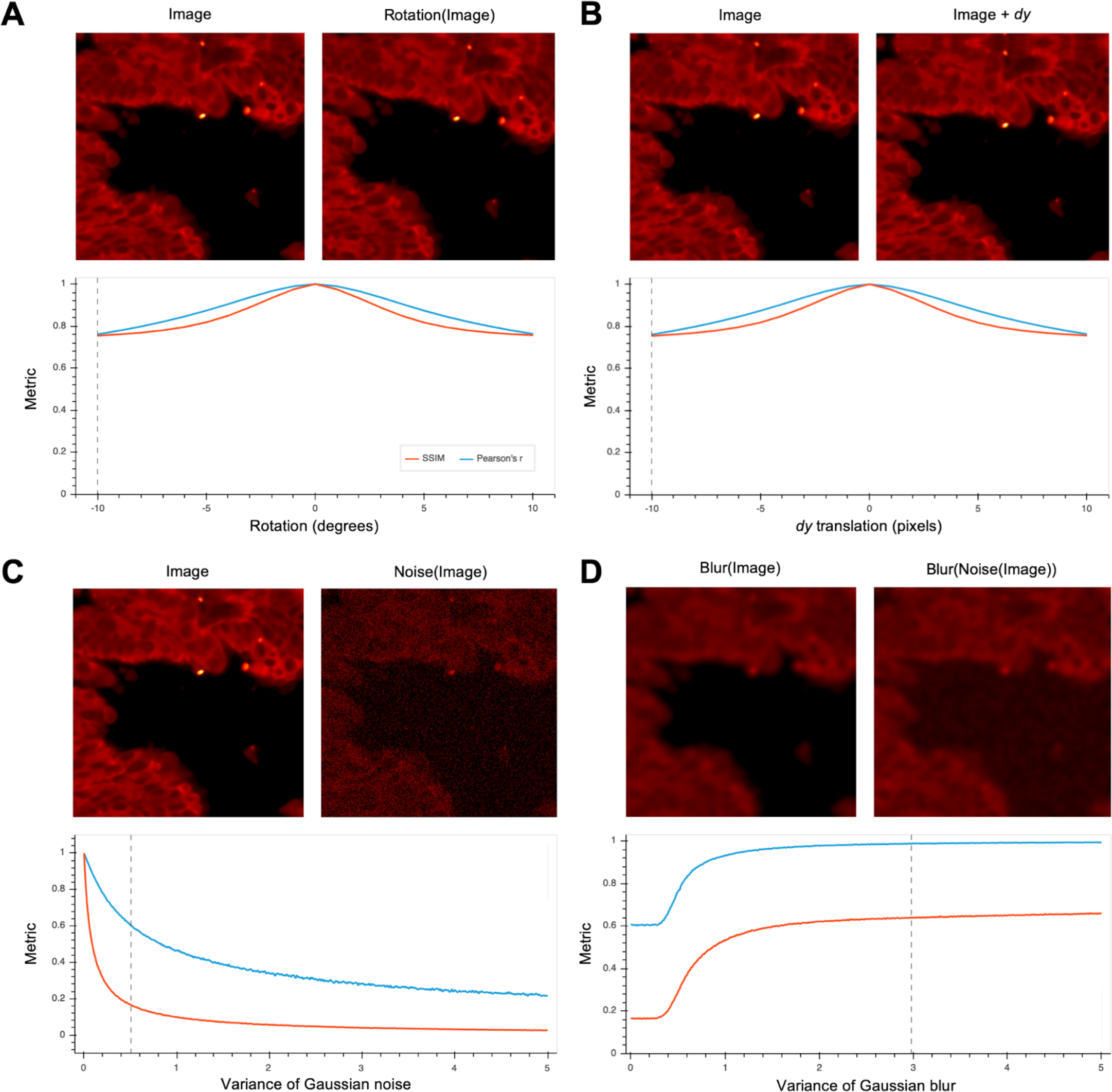
Metric sensitivity to perturbations. Structural similarity (SSIM) and Pearson’s *r* are two commonly used metrics for image comparison, both having been used to make comparisons between real and generated biological images in recent related work (Christiansen et al., 2018; Ounkomol et al., 2018; Rivenson et al., 2019b). When considering a stereotypical panCK IF tile, both measures are found to be sensitive to rotation (A) and translation (B), perturbations common to image registration. Grey dotted lines indicate the parameter selected to generate each transformed image. We also observe that both measures are sensitive to simulations of technical noise (C). By applying a Gaussian filter with variance (sigma) set to 3, we recover the SSIM between real and perturbed IF images without sacrificing global image details.

